# Effects of transcranial photobiomodulation on peripheral biomarkers associated with oxidative stress and complex IV activity in the prefrontal cortex in rats subjected to chronic mild stress

**DOI:** 10.1101/2024.06.27.597940

**Authors:** Luciana Bortoluzzi, Rafael Colombo, Karoline Motta Pinto, Lucas Henriques Viscardi, Ricardo Missiaggia, Douglas Jean Turella, Lisandra Schwantess, Mirian Salvador, Catia Santos Branco, Marina Rigotti, Ellen Scotton, Tainá Schons, Silene Bazi Ribeiro, Marco Antonio Caldieraro, Adriane R Rosa

## Abstract

**Background:** This study addresses the urgent need for effective alternatives to treat major depressive disorder (MDD) in patients who do not respond to conventional therapies Transcranial photobiomodulation therapy (tPBM) shows promise by enhancing mitochondrial function and reducing oxidative stress, as demonstrated in the chronic mild stress (CMS) rat model.

To analyze the impact of tPBM with two wavelengths (red and infrared) on behavioral and biological parameters related to MDD in a CMS model.

**Methods:** Male Wistar rats were subjected to CMS for five weeks and categorized into resilient (CMS-R) and susceptible (CMS-S) groups using the sucrose consumption test (SCT). The CMS-S group received tPBM treatment (600nm and 840nm) for five weeks. Biological measures included lipid damage (TBARS), antioxidant defense (TEAC), mitochondrial complex IV activity (CCO), and nitric oxide (NO) concentration in the prefrontal cortex and blood were measured.

**Results:** As expected, post-tPBM treatment (both red and infrared groups) exhibited increased sucrose consumption compared to the sham (Kruskal Wallis chi-squared=26.131; p<0.001). The red and infrared presented higher serum TEAC levels than the sham and control groups, but these effects did not reach statistical significance (p=0.306). In contrast, the red group showed lower peripheral TBARS levels (M = 9.50, SD = 2.87) than the sham group (M = 13.66, SD = 2.20) (p=0.0048); such effect was similar to the control non-stress group. The infrared group showed higher NO levels within the hippocampus than the sham group (Mean = 107.83; SD = 6.48, Dunn Test p = 0.0134) and higher prefrontal CCO activity levels than the red group (p=0.012), which was similar to the control non-stress group.

**Conclusions:** Our study demonstrated that tPBM using both red and infrared wavelengths significantly improved behavioral and biological parameters in the chronic mild stress (CMS) rat model. In particular, tPBM may offer therapeutic benefits by ameliorating oxidative stress and enhancing mitochondrial function, thereby presenting a promising alternative for the management of MDD.

## 1. Introduction

Major depressive disorder (MDD) is a complex and multifaceted mental illness that presents with ongoing feelings of sadness, lack of interest in previously enjoyed activities, repetitive thoughts of death, and cognitive and physical symptoms (Dadkhah et al., 2023). According to the Global Burden of Disease Collaborative Network, MDD impacts 185 million individuals worldwide (Marx et al., 2023). People with MDD often present impairment in occupational, familial, and social domains, concomitantly diminishing the overall quality of life (Malhi and Mann, 2018).

MDD is often accompanied by physical comorbidities, which may contribute to a decreased life expectancy in individuals with MDD (Arnaud et al., 2022). Indeed, depressive symptomatology carries consequences such as an increased incidence of cardiovascular diseases (Rajan et al., 2020), including coronary artery disease, heart failure, and stroke (Silva et al., 2017). Further, MDD increases the risk of suicidal ideation, attempts, and death, contributing to its disease burden (Su et al., 2023).

The standard treatment recommendations typically encompass a combination of antidepressant pharmacotherapy and psychotherapeutic interventions (Marx et al., 2023). However, it is noteworthy that a substantial proportion, approximately one-third of individuals diagnosed with MDD, fail to attain a satisfactory clinical response despite undergoing multiple therapeutic interventions (McAllister-Williams et al., 2020). Consequently, such cases are classified as treatment-resistant depression (TRD) (Amasi-Hartoonian et al., 2022). Consequently, the importance of identifying and implementing effective adjuvant therapeutic interventions in managing TRD becomes patently evident.

Chronic Mild Stress (CMS), first described almost four decades ago, has become a popular model in developing drugs for treating MDD. This model has engendered a substantial number of scientific investigations, with nearly 1700 studies solely focusing on rat-based research (Strekalova et al., 2022). Chronic stress exerts many harmful effects, encompassing mood, cognitive, and memory impairments. Research has shown that unpredictable stress can seriously affect the brain, including atrophy in the cortical and limbic regions, sensitization of the serotonergic system, increased levels of inflammatory proteins in the hippocampus, proliferation, and microglial activation, and disruptions in the HPA axis (Antoniuk et al., 2019). Moreover, this model has been extensively used to mimic animal models of depression.

Current research on mood disorders has focused heavily on the intricate relationship between systemic inflammation, oxidative stress, and the brain. The disparity between the production of reactive oxygen species (ROS) and the body’s capacity to detoxify them, commonly referred to as oxidative stress, has been associated with the emergence of MDD. This disparity can damage neuronal structures and neurotransmitter systems, which can contribute to the onset and persistence of depression (Gałecki and Talarowska, 2018). Emerging evidence suggests that oxidative stress and systemic inflammation may synergistically exacerbate depressive symptoms, highlighting the intricate web of factors involved in the pathogenesis of MDD and offering potential targets for innovative therapeutic interventions (Beurel et al., 2020).

Photobiomodulation (PBM) emerged as a promising treatment for MDD (Caldieraro and Cassano, 2019). This therapy uses red to near-infrared (NIR) spectrum photons (600-1100 nm) to irradiate tissue (Salehpour et al., 2020); using lasers or light-emitting diodes (LEDs) as the light source. The transcranial photobiomodulation modality (tPBM) is a novel therapy that employs visible or non-visible light delivered to the skull aiming to modulate the activity of the subjacent cortex. Studies have demonstrated that red and near-infrared light with wavelengths ranging from 630 to 810 nm can penetrate the skull at a rate of 0.2% to 10% (Salehpour et al., 2020). Light in the red and NIR spectrum is absorbed by the cytochrome C oxidase (CCO), triggering a series of cellular and physiological events, such as stimulating mitochondrial metabolism, increasing adenosine triphosphate (ATP) production, decreasing oxidative stress, neuroinflammation, and neuronal apoptosis, repairing damaged hypoxic cells, modulation of target tissue function and augmented cerebral metabolism and perfusion by releasing nitric oxide (NO) (Cassano et al., 2015; Montazeri et al., 2022).

Additionally, tPBM has been utilized to treat mood disorders in both animal models and humans and has displayed the potential to alter the neuroinflammatory profile in the brains of young and old rats (Cardoso et al., 2022; Montazeri et al., 2022). Furthermore, tPBM is considered safe when delivered by trained professionals or self-administered at home, depending on the device used for the treatment (Cassano et al., 2015). It typically does not cause pain, significant heat, or tissue damage, making it a non-invasive and low-risk therapy (Hamblin, 2017).

Although t-PBM is a promising treatment for MDD, the ideal stimulation parameters, including the most effective wavelengths, are not yet established. Also, the mechanisms of its antidepressant effect are not completely understood. Therefore, this study aimed to analyze the impact of tPBM with two different wavelengths (red 600 nm and NIR 840 nm) on behavioral, peripheral, and central biological parameters related to the pathophysiology of MDD in a CMS model.

## 2. Materials and Methods

### 2.1 Animals

We used eighty-eight (N = 88) male Wistar rats from the Laboratory Animal Reproduction and Experimentation Center of the Federal University of Rio Grande do Sul (CREAL-UFRGS). The animals were two months old and weighed approximately 250 - 300g. To calculate the sample size, we used the mean and standard deviation data for the sucrose consumption outcome of the control groups and those submitted to the CMS protocol, evaluated in the study by Wang and collaborators (2014) (Wang et al., 2014). We used the ClinCalc system, available at (https://clincalc.com/Stats/SampleSize.aspx) for the calculation. We considered the difference between the means and standard deviations for each calculated outcome, an alpha error probability of 0.05 (p<0.05), and a confidence interval of 90%. The estimated value was ten animals per group. However, based on the literature, it was considered that only 50% of animals subjected to stress would develop depressive-like behavior (Golden et al., 2011; Nieto-Gonzalez et al., 2015). Thus, to apply the CMS protocol and categorize groups as susceptible or resilient, a duplicated initial “N” is required. Therefore, 88 animals were needed (n = 4 in the pilot study; n = 14 animals excluded due to variation in sucrose consumption; n = 10 in the control group; n = 60 in the CMS group).

The animals were housed in the Physiology and Pharmacology Laboratory of the University of Caxias do Sul (UCS) under standard conditions of a 12-hour light/dark cycle, temperature of 22 ± 2°C, relative air humidity of 40 - 60%, with standard food and water *ad libitum.* Before beginning the CMS protocol, the animals were quarantined and acclimated for 14 days, with three animals per box. After that, basal sucrose consumption was measured. During the CMS protocol, animals were housed individually and kept isolated until the experiments ended. The animals in the control group were housed in groups (2 per box) to prevent them from suffering isolation stress. This study was approved by the Animal Ethics Committee (CEUA) of the University of Caxias do Sul under number #002/2020 on 03/13/2020.

### 2.2 Analysis of Light Penetration Energy Delivery

Before starting the experiment, a pilot study was carried out to evaluate the delivery of light energy and the penetration of these lights into the tissues, in addition to heating on the skin surface using red (600nm) and infrared (840nm) LED lights. Tissue samples were used from four animals (two animals aged two months and two animals aged four months) of the same species as those subjected to the study protocol. The samples comprised the frontal portion of the head (skull), skin, and hair. The samples were irradiated, two with each wavelength, at the same point on the skull where the animals received the treatment. The amount of energy that penetrated the samples was measured for each of the wavelengths: the bone part with scalp and hair and the bone part with scalp and without hair. Therefore, no significant heating of the skin surface was observed. The bone part with scalp and without hair had better light penetration, with 17%±6% of red light (600nm) and 19%±4% of infrared light (840nm) applied to the skin surface able to cross these structures when compared to the bony part with scalp and hair (9%+-3% of red light and 10%+-2% of infrared light. Table 1 describes the doses used in this study. The samples were measured using an optical sensor connected to a digital source (Icel Manaus - model PS-6000). The samples were measured using an optical sensor connected to a digital source (Icel Manaus - model PS-6000). The surface temperature of the sample was detected by a thermographic camera (FLIR System AB). A digital caliper (MTX-316119) was used to measure the thickness of the irradiated bone region. The LED on the sensor was held in position for approximately 10 seconds, or until the penetration power readings stagnated at a maximum of 20s, to obtain penetration readings (mW). This procedure was carried out in duplicate for each evaluation by the engineering sector of the company Tonederm®, which also provided the LED prototypes for application in this experiment.

**Table 1.** Sucrose consumption (g) during stress.

|  | Baseline | Week 2 | Week 3 | Week 4 | Week 5 | Week 5/Baseline |
| --- | --- | --- | --- | --- | --- | --- |
|  | *M (SD)* |  |  |  |  |  |
| **Control** | 3.04 (3.38) | 5.93 (4.53) | 14.33<br>(23.98) | 6.89 (5.77) | 5.05 (4.14) | 1.661 |
| **Infrared** | 5.67 (3.26) | 4.42 (7.39) | 10.61 (24.2) | 4.30 (2.69) | 1.81 (1.35) | 0.319 |
| **Red** | 3.40 (4.01) | 3.95 (7.01) | 11.1 (28.88) | 2.69(1.48) | 1.83(1.32) | 0.538 |
| **Sham** | 3.23 (2.29) | 12.74<br>(31.07) | 3.05 (2.67) | 5.26(6.48) | 1.80 (1.37) | 0.557 |
| **chi-squared** | 5.7761 | 3.3178 | 2.3691 | 3.5927 | 7.5069 |  |
| **p-value** | 0.123 | 0.345 | 0.499 | 0.309 | **0.0574** |  |

**Table 2:**
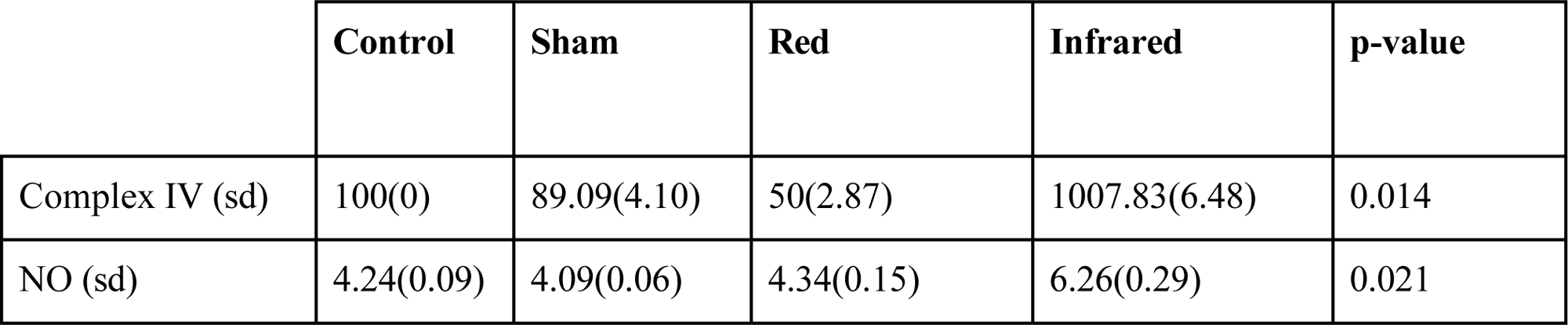
Nitric Oxide (NO) and Complex IV Values.

### 2.3 Sucrose consumption - Baseline

After two weeks of quarantine and acclimatization, the rats were introduced to sucrose consumption. The animals were trained to consume a 1% sucrose solution twice a week for 30 days. After 14 hours of food and water deprivation, the animals (now individualized) received a freshly prepared 1% sucrose solution for 60 minutes in a 100ml bottle numbered according to each animal. Sucrose intake was calculated by weighing the bottles before and after the test. The basal sucrose consumption of each animal was evaluated in the last week.

Papp (2012) found that animals showed variation in sucrose consumption before starting the stress protocol(Papp and Willner, 2023). This significant variation may affect our results, as the sucrose test is the primary outcome in characterizing anhedonic behavior. This study demonstrated that 20% of the animals present either a high consumption, a reduced consumption, or a significant variation in sucrose consumption. Animals that fit this profile were excluded from the experiment (n=14).

To divide the animals into stressed and control groups, they were paired based on their average sucrose consumption. Sucrose consumption was monitored weekly under similar conditions throughout the CMS protocol until the end of the study. After the test, food and water were replaced after a one-hour wait.

### 2.4 Chronic Mild Stress Protocol (CMS)

The CMS protocol lasted ten weeks. Animals in the CMS group were exposed to stressors according to the method described by Papp et al. (2018) and Willner et al. (2019) (Papp et al., 2018; Willner et al., 2019). The rats belonging to the CMS group were placed in standard plastic boxes and kept separate from each other. They were exposed to the following stressors: two (02) weekly periods of food and water deprivation; two (02) weekly periods with the cage tilted at 45°; two (02) weekly periods of intermittent lighting on/off every two (02) hours; two (02) weekly periods with a dirty cage; two (02) weekly periods with the presence of an intruding animal; two (02) weekly periods with low-intensity stroboscopic lighting; two (02) weekly stress-free periods. For ten consecutive weeks, stress was induced once or twice daily (10-14 hours for each stressor).

After conducting the sucrose consumption test in the fifth week of the CMS protocol, animals were divided into susceptible and resilient subgroups. The susceptible subgroup (CMS-S) consisted of animals that exhibited depressive-like behavior after the behavioral protocols. In contrast, the resilient subgroup (CMS-R) included animals that did not display such behavior. The definition of susceptible or resilient animals was guided by the outcome observed in the sucrose consumption test, defining susceptible animals as those with sucrose consumption lower than 40% of the baseline assessment value (Papp, 2012).

After 5 weeks of CMS, fifty percent of the animals were classified as susceptible to stress; treatment with tPBM red (600nm) or tPBM infrared (840nm) was started for five weeks. The animals in the CMS-R group were euthanized, while the CMS-S group continued to be subjected to stress and treatment with tPBM. The CMS-S group received treatment using tPBM, with each group having similar sucrose consumption averages through a pairing considering the sucrose consumption outcome. This led to the group’s division into three new subgroups based on the wavelengths of 600nm (Red group), 840nm (Infrared group), and a sham group. The animals in the non-stress control group were handled daily for the same period as those in the stress group.

### 2.5 tPBM treatment

Treatment with both wavelengths was transcranial. The animals were manually immobilized, and the light source was positioned in the frontal portion of the animal’s head (dorsal midline of the frontal region), aiming to stimulate the prefrontal cortex (Salehpour et al., 2018). Table 1 describes the parameters used. All animals underwent trichotomy at the application site, as the hair makes it difficult for light to pass through. Treatment with transcranial photobiomodulation occurred once a day for five weeks.

**Table 1:** Parameters used in transcranial photobiomodulation treatment.

| Parameter | Value | Value |
| --- | --- | --- |
| Wavelength | 600nm | 840nm |
| Light source | LED | LED |
| Wave mode | continuous | continuous |
| Illumination time | 150 seconds | 150 seconds |
| Spot size | 0,25 cm<sup>2</sup> | 0,25 cm<sup>2</sup> |
| Total power output | 400 mW | 400 mW |
| Total output power density | 1,6 W/cm<sup>2</sup> | 1,6 W/cm<sup>2</sup> |
| Average light penetration in the PFC | 17% ± 6% | 19% ± 4% |
| Estimated total delivery power in the PFC | 68 mW ± 4,0 | 76 mW ± 3,0% |
| Estimated power density in the PFC | 272 mW/cm<sup>2</sup> ± 16,3 | 304 mW/cm<sup>2</sup> ± 12,1 |
| Total energy per session | 10,2 J ± 0,6 | 11,4 J ± 0,4 |
| Estimated fluency delivered to the PFC | 40 J/cm<sup>2</sup> ± 2,4 | 45 J/cm<sup>2</sup> ± 1,8 |

### 2.6 Euthanasia and collection of biological material

Twenty-four hours after the last sucrose consumption test, the animals were euthanized. Animal euthanasia followed the procedures and methods indicated by RN 37/2018 of the CONCEA Euthanasia Practice Guidelines and described in project n° 15-0353, previously approved. The method used was anesthetic with intranasal isoflurane overdose followed by decapitation. The brain was immediately removed and dissected, separating the region of the prefrontal cortex and the hippocampus. The truncal blood was drawn shortly after the decapitation. The tissues were frozen in liquid nitrogen for subsequent storage in an ultra-freezer (−80°C) and biochemical evaluation.

### 2.7 Measures of lipid damage and antioxidant defense on peripheral blood

#### 2.7.1 Thiobarbituric acid reactive substances (TBARS)

Lipid peroxidation was monitored by the formation of thiobarbituric acid reactive substances (TBARS) during an acid-heating reaction, according to Wills (1966), with modifications (Wills, 1966). Specifically, 400 µL of supernatant from each sample was combined with 600 µL of 15% trichloroacetic acid and 0,67% thiobarbituric acid. The mixture was heated at 100°C for 15 min. After cooling to room temperature, the samples were centrifuged at 5200 xg for 30 min. The supernatants were isolated, and their absorbance was measured at 530 nm. Hydrolyzed malondialdehyde (MDA) was used as standard, and the results were expressed as nmol MDA/mL (Wills, 1966). The tests were conducted at the Oxidative Stress and Antioxidants Laboratory at the University of Caxias do Sul/RS - Brazil.

#### 2.7.2 Determination of Total Antioxidant Capacity Serum (TEAC)

The screening of antioxidant activity was performed through the sample’s ability to scavenge the radical ABTS+ [2,2-azino-bis (3-etilbenzotiazolin)-6-sulfonic acid]. The method follows the procedures described by (Re et al., 1999). The ABTS+ solution is formed from the reaction of 7 mM ABTS with 2.45 mM potassium persulphate. This solution was kept in the dark at room temperature for 12-16 hours before use. Then, the solution ABTS • + was diluted with 5mM phosphate buffer saline (PBS pH 7.4) until an absorbance of 0.700 ± 0.35 at 734 nm. Then, 1.0 mL of ABTS • + diluted solution was added to 10 µL of sample. The absorbances were read precisely 6 minutes after the initial mixture. A standard curve was used with Trolox solution for quantification, and the results were expressed in mM of equivalents of Trolox (Re et al., 1999). The tests were conducted at the Oxidative Stress and Antioxidants Laboratory at the University of Caxias do Sul/RS - Brazil.

### 2.8 Central measurements on cerebral tissue

#### 2.8.1 Complex IV activity in the prefrontal cortex

The activity of the mitochondrial chain complex IV, also known as cytochrome C oxidase (CCO), was assayed in the homogenized cerebral cortex (Spinazzi et al., 2012). The analyses were performed on all samples; however, due to the characteristics of the technique and the tissue under study, the samples were prepared in 3 pools per group. Complex IV activity was measured at 550 nm, and the positive control was potassium cyanide. Enzymatic activity was evaluated at 37°C in a spectrophotometer (Shimadzu model UV-1700). The results were presented as percentages. The tests were conducted at the Oxidative Stress and Antioxidants Laboratory at the University of Caxias do Sul/RS - Brazil.

#### 2.8.2 Nitric Oxide Concentration in the Hippocampus

Nitric oxide (NO) bioavailability was assessed according to the Griess reaction based on the Green et al. (1982) method (Green et al., 1982). Due to the characteristics of the technique and the tissue under study, the samples were prepared in 3 pools per group. The homogenized tissues were added to Griess reagent (1:1), mixed, and incubated in the dark for 10 min after reading at 550 nm by a microplate reader. Sodium nitroprusside was used as the standard. The results were expressed as nmol of nitrite/mg of protein. The tests were conducted at the Oxidative Stress and Antioxidants Laboratory at the University of Caxias do Sul/RS - Brazil.

### 2.9 Statistical Analysis

We analyzed the data using the R platform with Dplyr and broom packages. The Shapiro-Wilk test for normality was used to verify whether each parameter followed a normal distribution before comparing groups. The analysis of variance (ANOVA) was applied to parametric data. At the same time, the Kruskal-Wallis was used to compare groups for non-parametric data, followed by Bonferronís pairwise grouping with a significance level of p<0.05. The graphs were generated on *ggpubr* and *ggplot2*, also within the R platform.

## 3. RESULTS

### 3.1 Sucrose consumption

#### 3.1.1 Baseline

The groups did not differ significantly at the baseline before the chronic stress protocol (Kruskal-Wallis, p=0.123). By the fifth week of protocol, the sham, red, and infrared groups showed lower sucrose consumption levels than the control group, without a statistical difference (Kruskal-Wallis chi-squared 7.5069; p=0.0574). On the first day of the tPBM intervention, the control group consumed 5,396g of sucrose, whereas the rats exposed to the stress protocol consumed 1,478g (sham) (p=0.00879). Indeed, the sucrose consumption was reduced by over 55% and 67% in the sham group and red/infrared groups, respectively, when compared to baseline measurements (Table 1).

#### 3.1.2 After transcranial Photobiomodulation

After five weeks of CMS, CMS-S rats received t-PBM for five weeks in addition to maintaining the CMS protocol for the same period. On the fifth week of treatment, both red and infrared groups exhibited increased sucrose consumption compared to the sham (Kruskal Wallis chi-squared=26.131; p<0.001). While the sham group remained in an anhedonic state, experiencing a 25% reduction in sucrose consumption, the red group more than doubled their consumption compared to the first day of the intervention. The infrared group recovered approximately 1,67 times their initial consumption level to the baseline. These data suggest that the tPBM using both infrared and red frequencies improved the anhedonic behavior in rats exposed to chronic stress protocol, as shown in Figure 1. It is important to highlight that this difference concerning initial consumption in the treated groups (red and infrared) does not present a statistical difference compared to the sham group. Furthermore, after 5 weeks of treatment with photobiomodulation, the infrared group showed no statistical difference compared to the control group.

**Figure 1.**
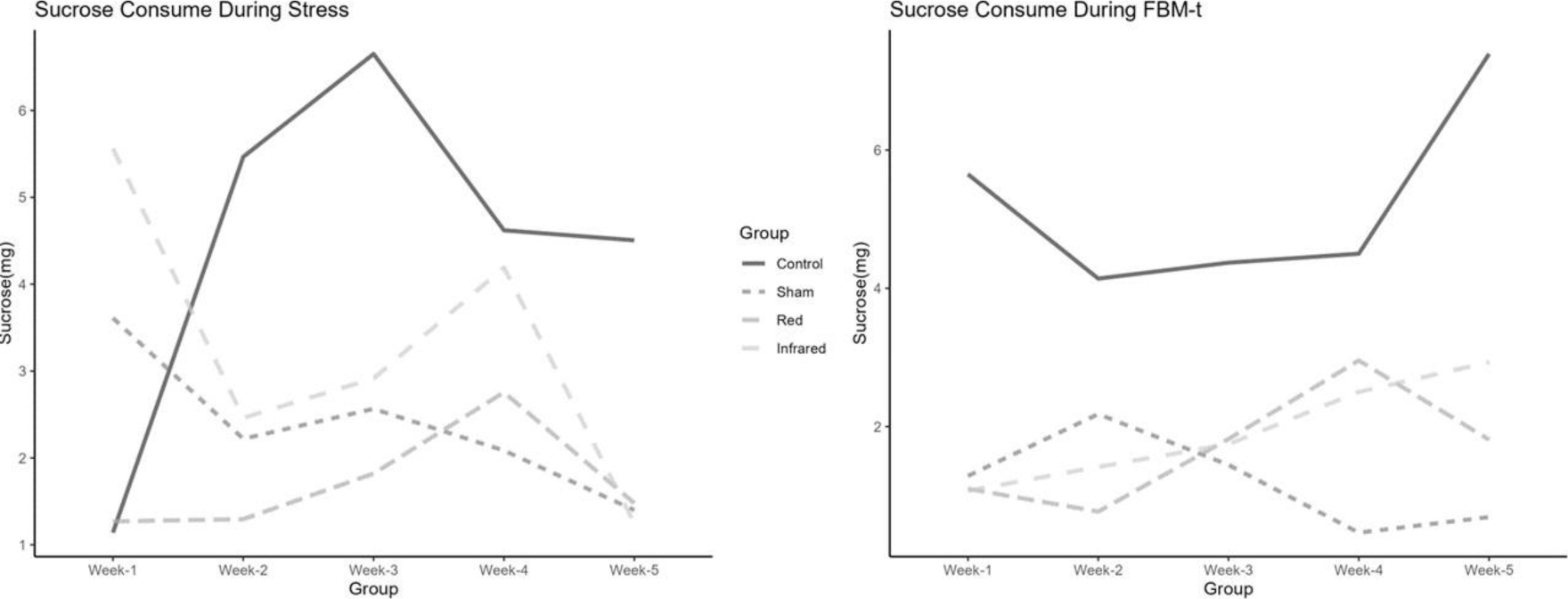
Sucrose consumption during chronic exposure to stress protocol and Transcranial Photobiomodulation treatment.

### 3.2 Peripheral blood biomarkers

#### 3.2.1 Thiobarbituric acid reactive substances (TBARS)

One-way ANOVA revealed statistical differences in TBARS levels among groups (F=3.21; p=0.0394). Post hoc analysis indicated that the red group showed lower TBARS levels (M = 9.50, SD = 2.87) than the sham group (M = 13.66, SD = 2.20) (p=0.0048) and was similar to the control group. There were no differences between infrared and other groups.

**Figure 2.**
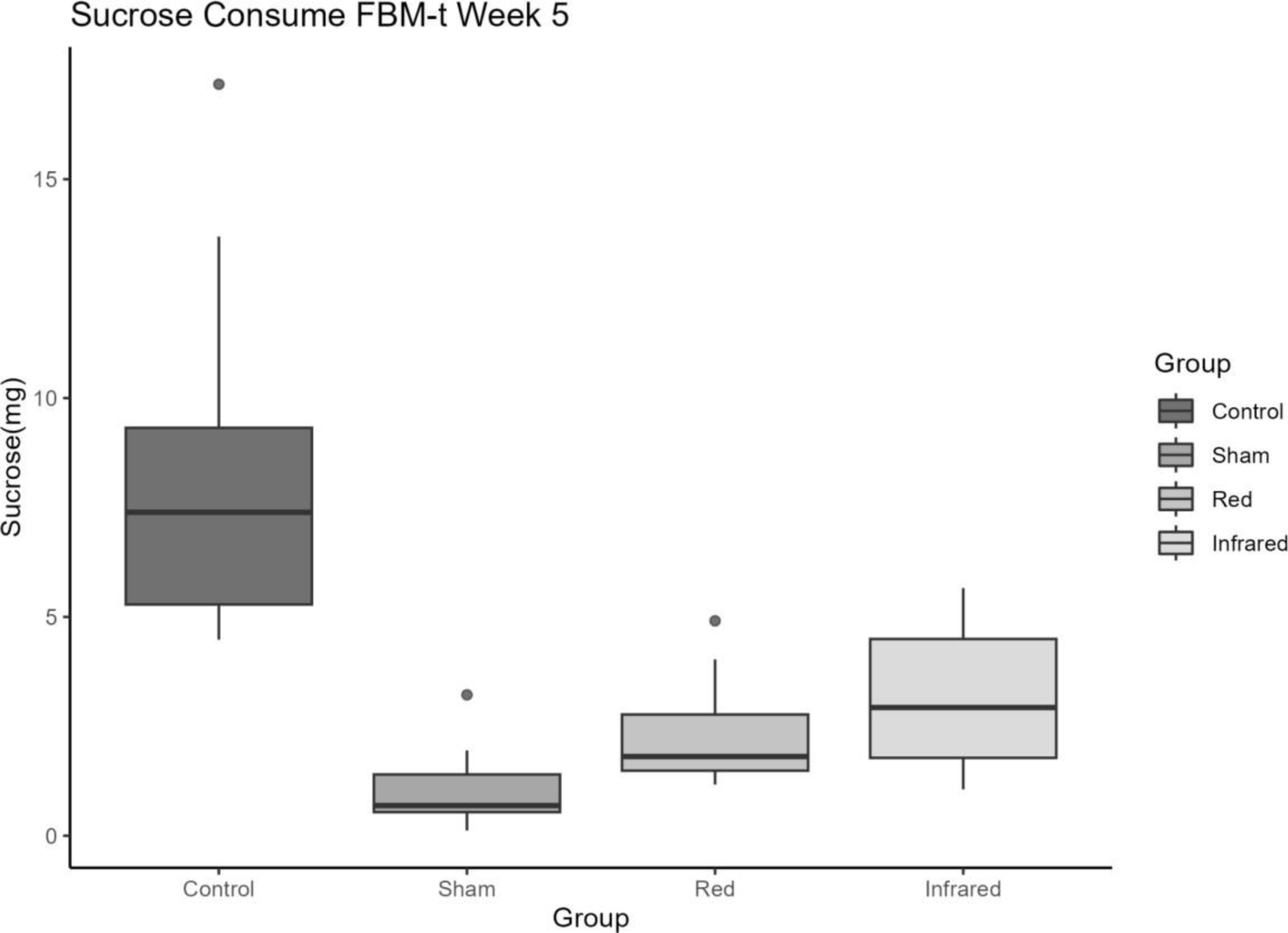
Sucrose consumption on the fifth week of chronic exposure to stress protocol and Transcranial Photobiomodulation treatment.

#### 3.2.2 Trolox equivalent antioxidant capacity (TEAC)

Although infrared and red groups presented higher TEAC levels than sham and control groups (Figure XX), these effects did not reach statistical significance (p=0.306).

**Figure 3.**
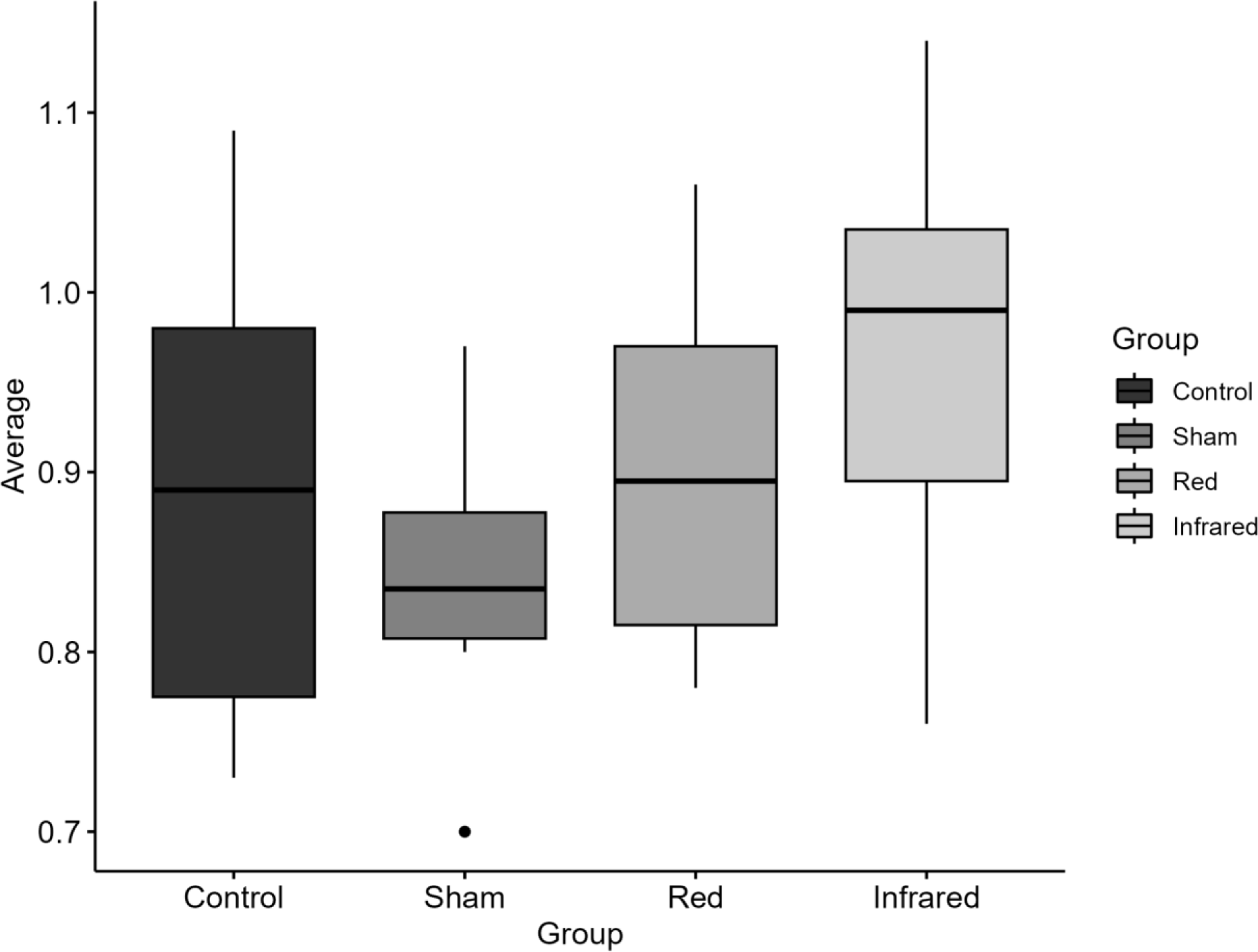
Comparison among values of Total Antioxidant Capacity Serum (TEAC).

**Figure 4.**
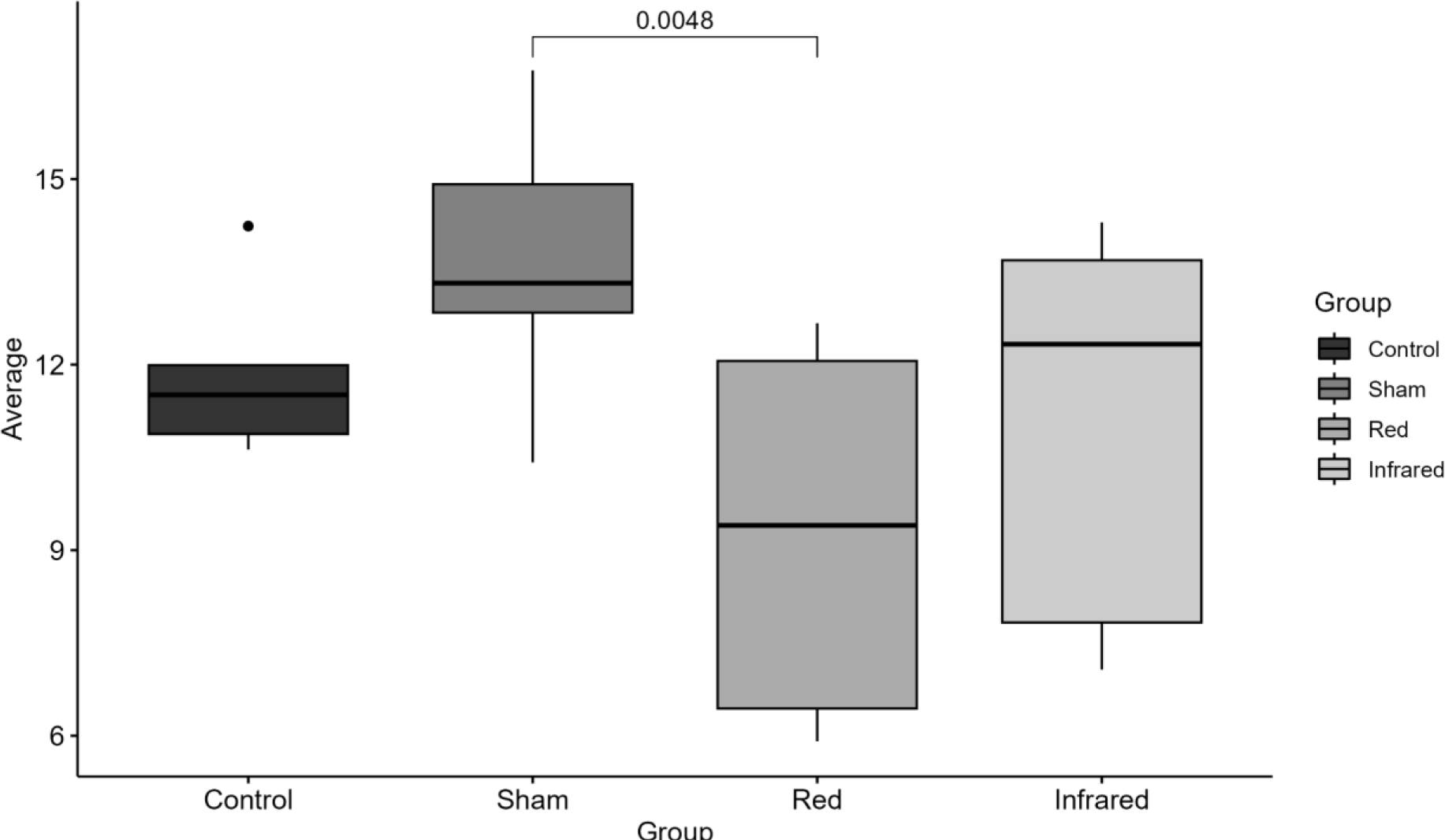
Comparison among values of thiobarbituric acid reactive substances (TBARS)

### 3.3 Central biomarkers

#### 3.3.1 Nitric oxide (NO) on the hippocampus

According to the Kruskal-Wallis test, there was a statistically significant difference in the hippocampal levels of NO among groups (Chi-squared = 9.6667; p=0.02162). In particular, the NO levels within the hippocampus of rats treated with infrared significantly increased compared to the Sham group, although not with the other two groups (Mean = 107.83; SD = 6.48, Dunn Test p = 0.0134). Conversely, no differences were observed between the red and other groups, as shown in Figure 5.

**Figure 5.**
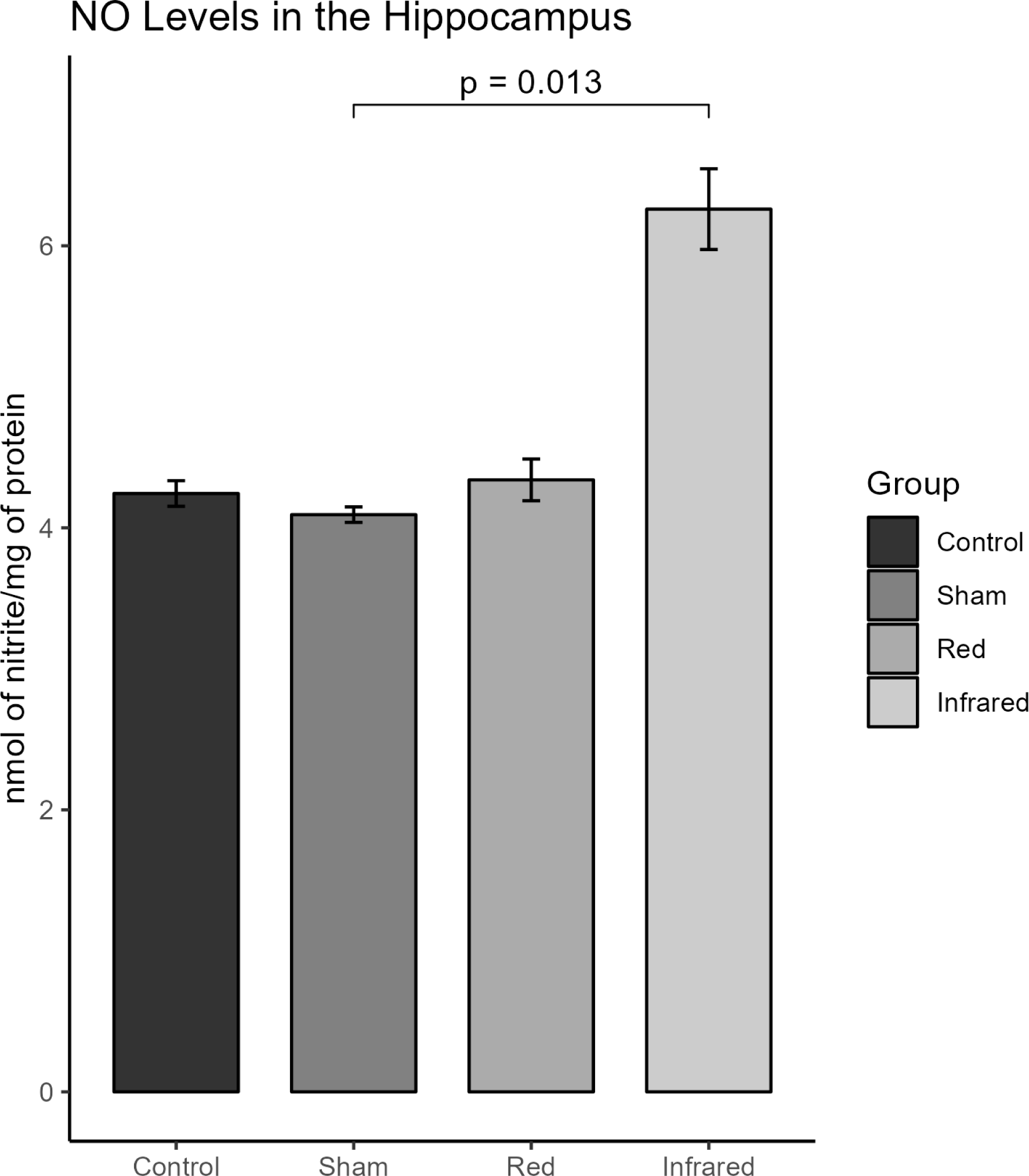
Effects of tPBM on hippocampal NO levels.

**Figure 6.**
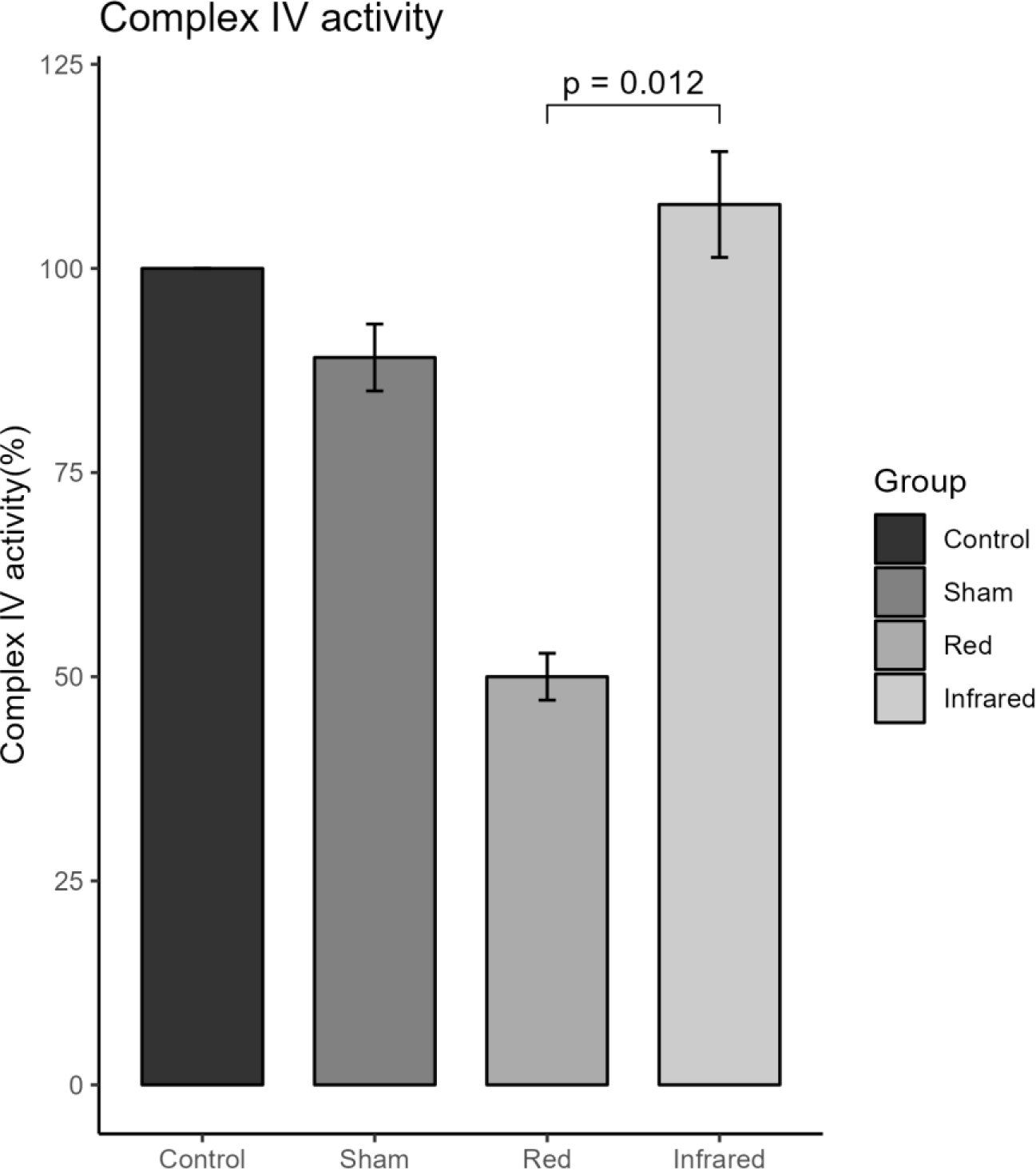
Mitochondrial complex IV activity of the cerebral cortex.

#### 3.3.2 Mitochondrial chain complex IV activity on the prefrontal cortex

The activity of mitochondrial complex IV (CCO) in the prefrontal cortex differed significantly among treatment groups (Kruskal-Wallis, Chi-squared = 10.532; p<0.01455). Post hoc analysis indicated that the infrared group exhibited higher CCO activity levels than the red (p=0.012), and it was similar to the control non-stress group. In contrast, the red group showed lower CCO levels than other groups (p<0.05).

## 4. Discussion

Our study showed that the CMS model efficiently reproduced anhedonic behavior showing a significant reduction of sucrose consumption in the groups exposed to this protocol. Furthermore, the tPBM red and infrared were able to reverse this depressive-like behavior, which suggests the promising antidepressant properties of this intervention. In the study by Xu et al. (2017), it was also shown that photobiomodulation effectively improved depression-like behaviors in two mouse models of depression(Xu et al., 2017). The CMS protocol is a well-established animal model for depressive-like behavior and is closely associated with anhedonic tendencies in rodents(Papp et al., 1994; Willner, 2017; Willner et al., 1987). Anhedonia is considered a central symptom in depressive disorders and also serves as a marker for predicting the prognosis of treatment-resistant depression(Coccurello, 2019; Scheggi et al., 2018; Stanton et al., 2019). Indeed, anhedonia has been used as a primary outcome of distinct CMS (Antoniuk et al., 2019). The sucrose preference test, initially proposed by Willner (1987) and posteriorly adapted by Papp (1994), stands as a gold standard for assessing anhedonia in rats(Papp et al., 1994; Willner, 2017; Willner et al., 1987). Our results showed the positive effects of tPBM in a valid animal model for depressive-like behavior, as observed in the study by Tanaka et al (2011)(Tanaka et al., 2011).

Several theories exist to explain the complex causes of MDD, including the inflammatory theory(Gałecki and Talarowska, 2018). Inflammation or the inflammatory response is a consequence of the activation of the immune system, typically appearing as a localized reaction. Its primary purpose is to rid the body of harmful stimuli or insults. Various immune cells and mechanisms are mobilized to safeguard the body’s internal equilibrium, known as homeostasis. In the brain, studies have shown that microglial activation leads to neuroinflammation in both animal models and clinically depressed patients(Wang et al., 2022). Thus, it is important to note that the dysregulation of the immune processes can frequently play a role in developing diseases, including MDD(Beurel et al., 2020). In line with this, oxidative stress (OS) is a biological phenomenon arising from the disruption in the balance between the production of reactive oxygen species (ROS), commonly known as free radicals, and the body’s ability to neutralize them through antioxidative defenses(Teleanu et al., 2022). The resultant impact of these ROS on cellular components can induce cellular dysfunction and trigger an inflammatory response, where inflammatory cells contribute to the generation of ROS and, consequently, OS(Lima et al., 2022; Teleanu et al., 2022). This connection reinforces the significant role of inflammation in mood disorders. Individuals with mood disturbances often display heightened levels of pro-inflammatory cytokines, oxidative stress products, chemokines, and soluble adhesion molecules in blood and cerebrospinal fluid(Wei et al., 2022). Furthermore, systemic inflammation and OS can contribute to neuroinflammation, a well-established factor in developing neurodegenerative diseases and mood disorders(Teleanu et al., 2022).

The present study investigated the antioxidant and metabolic properties of tPBM using the TBARS, TEAC, NO, and mitochondrial complex on cortical structures and serum of rats exposed to CMS protocol. Both red and infrared wavelengths produced lower TBARS levels when compared to sham stimulation, although only the red group demonstrated statistical significance. Regarding TEAC levels, both infrared and red groups exhibited higher TEAC levels compared to the sham and control groups. These findings indicated that tPBM, particularly the red group, showed a better profile of oxidative lipid damage, even without statistical difference. Even with the absence of a statistical difference in the measurement of antioxidant capacity, the reduction seen in this outcome may indicate a biological effect of tPBM on the oxidative stress profile. In this context, the study by Yang et al. (2011) demonstrated the ability of red light (632.8 nm) to suppress Aβ-induced ROS production and inflammatory response in rat primary astrocytes (Yang et al., 2010). A study using a restraint stress model mice model showed that tPBM with infrared light (810nm) and Coenzyme Q10 improved the antioxidant defense capacity of the brain(Salehpour et al., 2019). In another study, Salehpour and colleagues showed that tPBM with infrared light reduced oxidative damage in the hippocampus, demonstrating the ability of 810nm laser light to enhance the antioxidant defense system and maintain mitochondrial survival in sleep-deprived (SD) mice. Treatment with transcranial NIR laser in SD mice restored hippocampal total antioxidant capacity levels compared to those in the untreated SD group(Salehpour et al., 2018). Together, these findings suggest that tPBM provides systemic and central nervous system antioxidant properties, suggesting the neuroprotective function of tPBM(Lu et al., 2023).

Furthermore, our results demonstrated that tPMB, particularly infrared, improved CCO (mitochondrial chain complex IV) activity significantly more than sham, achieving levels of activity similar to those observed in the control non-stressed group. The mitochondrial complexes are sets of enzymatic proteins located in the inner membrane of mitochondria. These complexes are crucial in the respiratory chain, the central metabolic pathway for ATP generation. Four main complexes (I, II, III, and IV) comprise multiple protein subunits collaborating in electron transfer along the respiratory chain. Mitochondrial complexes are pivotal for cellular energy homeostasis, playing a fundamental role in the healthy functioning of cells and organisms(Sousa et al., 2018). Absorption of photon energy by the CCO is proposed to be a central mechanism for the PBM effect(Hamblin, 2016). For instance, laser treatment using both red and infrared (660 and 810 nm) has shown decreased reactive oxygen species (ROS) production and increased cytochrome c oxidase (CCO) compared to sham in the D-Galactose-induced aging model(Salehpour et al., 2017). Another research identified changes in CCO activity within cortical and subcortical regions after tPBM with an infrared laser administration (1,064nm)(Wade et al., 2023).

The study by Wade et al. (2023) reports that the most significant shifts in CCO activity occurred one day after tPBM with infrared light application in the infralimbic prefrontal cortex, with effects persisting elevated for 2 to 4 weeks post-tPBM(Wade et al., 2023). Moreover, significant differences in CCO activity between the 2-week and sham groups were observed in various brain regions, including the molecular layer of the hippocampus, the CA3 region of the hippocampus, the lateral septum, and the nucleus accumbens. Interregional correlation analysis indicated enhanced functional connectivity between cortical and subcortical areas post-tPBM, which endured for four weeks post-stimulation. The temporal pattern of changes in CCO activity and functional coupling suggests the occurrence of distinct forms of neuroenergetic plasticity at different time intervals post-tPBM, contingent upon the brain region and its cortical depth.

In this context, evidence has shown that in vivo oxidation of CCO is a direct photonic action of administration of tPBM to the human prefrontal cortex. Upon penetrating the skull, the tPBM is absorbed by CCO, leading to a direct and non-thermal photonic oxidation of CCO(Wang et al., 2022). Wang et al. (2017) demonstrated that tPBM-NIR positively regulates oxidized cytochrome c oxidase (CCO) in the human brain, a finding corroborated by Pruitt et al. (2020), who also observed an increase in the concentration of oxidized CCO (Pruitt et al., 2020; Wang et al., 2017). Saucedo et al. (2021) advocate using tPBM to enhance cerebral oxygenation and alleviate age-related declines in mitochondrial respiration(Saucedo et al., 2021). They found significant elevations in ATP biosynthesis and the expression and activity levels of mitochondrial complex IV in the prefrontal cortex (PFC) following PBM. Additionally, Zhang et al. (2014) and Huang et al., (2013) have suggested the protective effects of PBM could be attributed to the increased ATP production and selective modulation of pro-inflammatory mediators(Huang et al., 2013; Zhang et al., 2014). Collectively, these studies underscore the impact of infrared tPBM on enhancing the bioenergetic capacity of the brain.

In our study, the t-PBM, particularly with NIR, increased nitric oxide (NO) in the hippocampus of treated animals. This may be correlated with the increased activity of CCO. This correlation arises from the fact that CCO can be inhibited by NO. PBM can dissociate NO from the CCO, leading to elevated mitochondrial membrane potential, increased oxygen consumption, enhanced glucose metabolism, and more outstanding ATP production by the mitochondria(Hamblin, 2017). It is important to highlight that the increase in NO concentration observed in the tPBM near-infrared group may be related to the increase in complex IV activity, resulting in its flow into the cytosol(Ghasemi et al., 2018). When released from mitochondrial regulatory sites, NO becomes available in the mitochondria and the cytosol, which may explain the higher NO levels observed in this group. Released NO can also act as a vasodilator, increasing blood flow and improving metabolism in the t-PBM irradiated area, also contributing to the antidepressant effect(Hamblin, 2016).

Regarding the limitations of this study, the appropriate dosage of tPMB for bio-stimulation induction is a challenge. This is because of factors such as wavelength, output power, continuous or pulsed emission, power density, irradiation time, dose in J/cm2, total delivered energy, application technique, and intervals between sessions(Dompe et al., 2020). Additionally, the three minutes of immobilization utilized in this experiment for tPBM application may have induced physical stress in the rats, potentially influencing the antidepressant effects of phototherapy. Furthermore, the CMS protocol has its own limitations, such as sex differences and interindividual vulnerability to stress, which are yet unexplored in this model(Loizeau et al., 2024).

## 5. Conclusion

In summary, our findings suggest that tPBM holds promise as an effective antidepressant strategy. The improvement in sucrose consumption, the increase in Complex IV activity in the prefrontal cortex, and the increase in NO concentration in the hippocampus, associated with a reduction in damage to membrane lipids peripherally, may be indicators of the beneficial effects of this therapy. For most of the outcomes assessed in this study, the NIR wavelength was superior or similar to red light, suggesting NIR should be preferred for future studies.

## Consent for publication

This study was approved by the University of Caxias do Sul’s Animal Ethics Committee (CEUA) on 03/13/2020 under number #002/2020.

## Data availability

The de-identified database used in the current study is available from the corresponding author upon reasonable request.

## Author contributions

Luciana Bortoluzzi: Conceptualization, Investigation, Writing - Original Draft. Rafael Colombo: Conceptualization, Methodology, Writing - Review & Editing, Project administration, Funding acquisition. Karol Motta Pinto: Investigation. Lucas Henriques Viscardi: Investigation, Formal analysis, Software. Ricardo Missiaggia:Investigation. Douglas Jean Turella: Investigation. Lisandra Schwantess: Investigation. Mirian Salvador: Investigation, Catia Santos Branco:Investigation. Marina Rigottib: Investigation. Tainá Schons: Writing - Original Draft, Visualization. Silene Bazi Ribeiro: Investigation. Marco Antonio Caldieraro: Writing - Review & Editing. Adriane Ribeiro Rosa: Writing - Review & Editing, Supervision, Project administration, Funding acquisition.

## Acknowledgments

Special thanks to: Angela Toniolli Padilha, Laura Bozzetti Suhnel, Ellen Scotton, and Guilherme Thomazi.

Luciana Bortoluzzi: Graduated in Physiotherapy and pursuing a Ph.D. in Pharmacology and Therapeutics. She specializes in dermatofunctional physiotherapy.

Rafael Colombo: He is an associate professor at Universidade de Caxias do Sul. His focuses on stress-related mood disorders and pharmacological resistance in major depressive disorder.

Holds a Ph.D. in Physiology, specializing in Cardiovascular and Neurophysiology. Lucas Henriques Viscardi: Background in Archaeology and Genetics. Research interests include human and non-human primate evolution, behavior genetics, and molecular evolution. He is also studying Medicine and is involved in HIV counseling research.

Mirian Salvador: Professor in Biochemistry, specializing in Oxidative Damage and Antioxidants. Actively publishes research papers and holds multiple patents in the field of natural products and redox metabolism.

Catia Santos Branco: Holds a Ph.D. in Biotechnology, focusing on oxidative stress and antioxidants. Leads research projects in cellular dynamics modulation and collaborates nationally and internationally. Also serves as an Associate Professor.

Ellen Scotton: She is biomedical scientist and PhD in Pharmacology and Therapeutics. Her focuses on tress-related mood disorders and pharmacological resistance in major depressive disorder.

Tainá Schons: Biomedical Scientist with expertise in bioinformatics and blood banking. Currently pursuing a PhD in Pharmacology focuses on molecular psychiatry and bioinformatics.

Silene Bazi Ribeiro: Physiotherapist specializing in Dermato-Functional Physiotherapy. Teaches in postgraduate courses and works as a scientific consultant for product research and development.

Marco Antonio Caldieraro: He works as Psychiatrist at the Hospital de Clínicas de Porto Alegre and is Professor of the Graduate Program in Psychiatry and Behavioral Sciences at the Federal University of Rio Grande do Sul (UFRGS). He holds a PhD in Medical Sciences and has completed Post-Doctoral studies on neuromodulation.

Adriane R Rosa: Associate Professor and Head of the Graduate Program in Pharmacology and Therapeutics at the Federal University of Rio Grande do Sul (UFRGS). She also coordinates the Laboratory of Molecular Psychiatry at the Hospital de Clínicas de Porto Alegre. Her focuses on the neurobiology of psychiatric disorders as well as the discovery of biomarkers and novel pharmacological targets. She holds a PhD in Medical Sciences and was fellow in the Neuroscience Institute at Hospital Clinic of Barcelona.

## Abbreviations

MDD: Major depressive disorde
TRD: Treatment-resistant depression
CMS: Chronic Mild Stress
ROS: Reactive oxygen species
PBM: Photobiomodulation
tPBM: Transcranial photobiomodulation
CCO: Cytochrome C oxidase
ATP: Adenosine triphosphate
NO: Nitric oxide
TBARS: Thiobarbituric acid reactive substances
TEAC: Total Antioxidant Capacity Serum

